# Room-temperature Storage of Lyophilized Engineered Bacteria using Tardigrade Intrinsically Disordered Proteins

**DOI:** 10.1101/2021.06.25.449888

**Authors:** Yixian Yang, Zhandong Jiao, Shao Zhang, Mingjian Shan, Sizhe Duan, Xinyuan Wang, Siyuan Wang, Yiming Tang, Shiqi Wang

**Affiliations:** Tsinghua University High School, Beijing 100084, China

**Author notes:** To whom correspondence should be addressed: Shiqi Wang.

**Keywords:** Tardigrade intrinsically disordered proteins, Freeze-drying resistance, Room temperature storage, Engineered bacteria

## Abstract

Tardigrades, which live in transiently wet environments such as moss, are well-known for their extreme resistance to desiccation. Tardigrade intrinsically disordered proteins (TDPs) have been reported to also protect bacteria and yeast under desiccation ^[4, 5, 32]^. In this study, we utilized lyophilization to achieve room-temperature storage of engineered bacteria. By using TDPs, engineered bacteria are protected under lyophilization and their original functions are preserved ^[12, 17, 18]^. This study shows that TDPs can be expressed in the *Escherichia coli* (*E. coli)* BL21 and DH5α, and bacteria treated with Cytosolic-abundant heat soluble protein (CAHS) 106094 displayed the highest survival rate after lyophilization ^[16, 41, 44]^. Moreover, this study shows that the co-expression of TDPs can improve the preservation of bacteria and maintain high survival rates after prolonged room temperature storage. Additionally, the TDPs can be expressed using different vectors, which means that they can be used in different types of engineered bacteria. This study offers a new storage method that not only improves the storage of biological material for industrial and daily usage, but also for future iGEM (International Genetically Engineered Machine Competition) teams to store and use their engineered bacteria in different applications.

## 1. Hypothesis

Trehalose is commonly used to protect diverse organisms and even enzymes ^[6, 10, 15, 28, 30, 34, 35]^ under conditions of desiccation ^[7, 26]^. However, it is not present in tardigrades, which are some of the most desiccation-resistant organisms on the planet ^[8, 14, 22, 33]^. Therefore, this study hypothesized that the tardigrade intrinsically disordered proteins (TDPs) ^[17, 18]^ are responsible for their desiccation resistance instead of trehalose ^[27]^. Since wild-type and engineered bacteria of the same species share similar structural features with the exception of edited genes, we hypothesized that TDPs can protect engineered bacteria during desiccation, as their ability to preserve common bacteria in a dry state was already demonstrated by Boothby and colleagues ^[4, 5]^. Additionally, this study further hypothesized that the lyophilization, or freezing and desiccation of engineered bacteria that are treated with TDPs would allow room-temperature preservation of engineered bacteria in the form of a dried powder that can be recovered and brought back to life as needed. Moreover, as different strains of *Escherichia coli* are structured similarly, this preservation method of engineered bacteria can be utilized widely for multiple strains of *E. coli*, and the combined expression of multiple TDPs can improve the survival rate after lyophilization.

## 2. Introduction

The lack of room temperature preservation of biological products is one of the most severe issues in current day biotechnology. Interviews done by QHFZ-China in 2020 revealed that 8 of the 10 interviewed iGEM teams believe that the issue of room temperature storage requires a solution. Moreover, other preservation methods for bacteria like the paraffin method are not all suitable for engineered bacteria and require trained staff, limiting their broader applications. By contrast, lyophilization is a viable option for room temperature preservation as it is easy to test on engineered bacteria and can be largely automated. However, although lyophilization can preserve substances in the form of a powder that can be stored at room temperature, research has shown that the survival rate of lyophilized of bacteria is less than 10%, meaning that most bacterial cells cannot survive the process. Luckily, there is a solution.

A study published in *Molecular Cell* by Thomas C. Boothby et al. in 2017 titled *Tardigrades Use Intrinsically Disordered Proteins to Survive Desiccation* tested the idea that tardigrade intrinsically disordered proteins (TDPs) can protect prokaryotic as well as eukaryotic cells and animals under conditions of desiccation ^[2, 4, 5, 14]^. The researchers discovered that heterologous expression of TDPs in *E. coli* BL21 (DE3) and yeast increased their ability to survive desiccation. This observation was mainly explained by the water replacement hypothesis. Unlike hypotheses that the molecules prevent cohesive interactions through interference ^[4, 25, 28, 37]^ or the vitrification hypothesis ^[9, 12]^, the water replacement hypothesis states that the hydrogen bond network of proteins is protected by TDPs, maintaining the normal structure of the protected protein in a dry environment ^[3]^. Using a process of TDP expression, lyophilization, and recovery of cells, this study aims to expand on the work done by Boothby et al., utilizing the protective effect of TDPs ^[20]^ for the preservation of engineered bacteria. This expansion of usage will allow longer preservation of pharmaceuticals that utilize engineered bacteria, preservation of crops produced with the help of engineered bacteria, as well as any products based on engineered bacteria such as vaccines, which require long-distance transportation or long-term storage without relying on an expensive, uninterrupted refrigeration chain.

## 3. Results

Result 1: TDPs can protect recombinant strains based on *E. coli* BL21 under conditions of lyophilization, allowing them to be preserved at room temperature ^[23]^. CAHS 106094 expression in BL21 resulted in the highest survival rate at room temperature after lyophilization.

**Figure 1.**
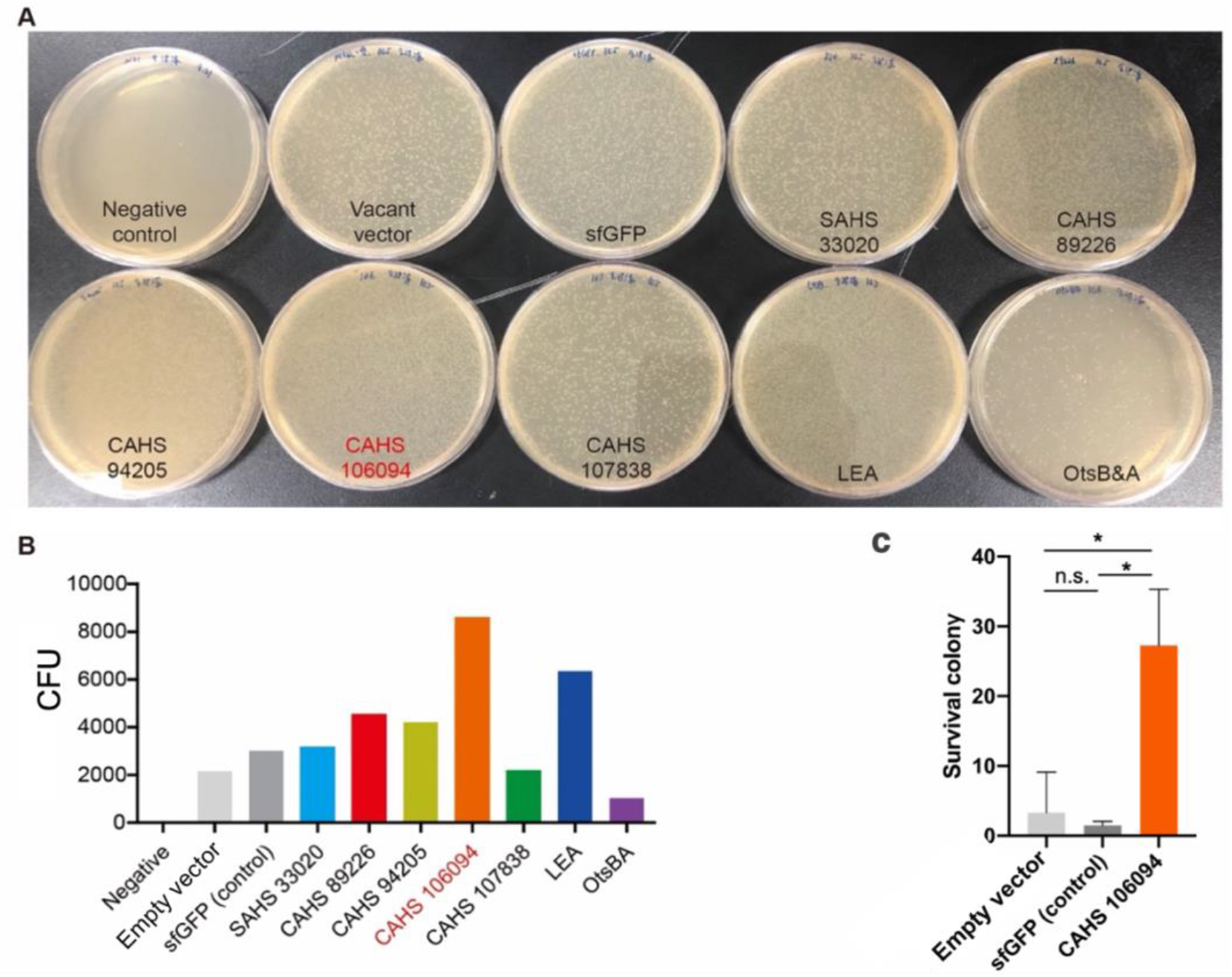
CAHS 106094 had the best protective effect against loss of viability during lyophilization. (A) Relative viability of *E. coli* BL21 strains expressing 5 different TDPs, LEA and OtsBA, respectively, streaked out on agar plates after lyophilization. (B) Quantification of the assay from (A), means ± SD from 2 technical replicates. The CFU value is calculated with a dilution factor of 10^5^. (C) Statistical analysis of the protective effect of CAHS 106094 compared to the control group. The data represent the means ± SD from 3 technical replicates.

*E. coli* BL21 was chosen because it was used in the desiccation study done by Boothby and colleagues ^[16]^. First, we utilized the T7 promoter, which has been proven to be the most efficient in the BL21 platform, and tested 4 CAHS proteins, 1 SAHS protein, and sfGFP as control ^[44]^. To select the optimal promoter for the expression of TDPs, we tested three constitutive promoters with varying strengths that are widely used in various strains of *E. coli*.

Result 2: Increasing the expression of the same TDP resulted in higher viability during storage at room temperature following lyophilization.

**Figure 2.**
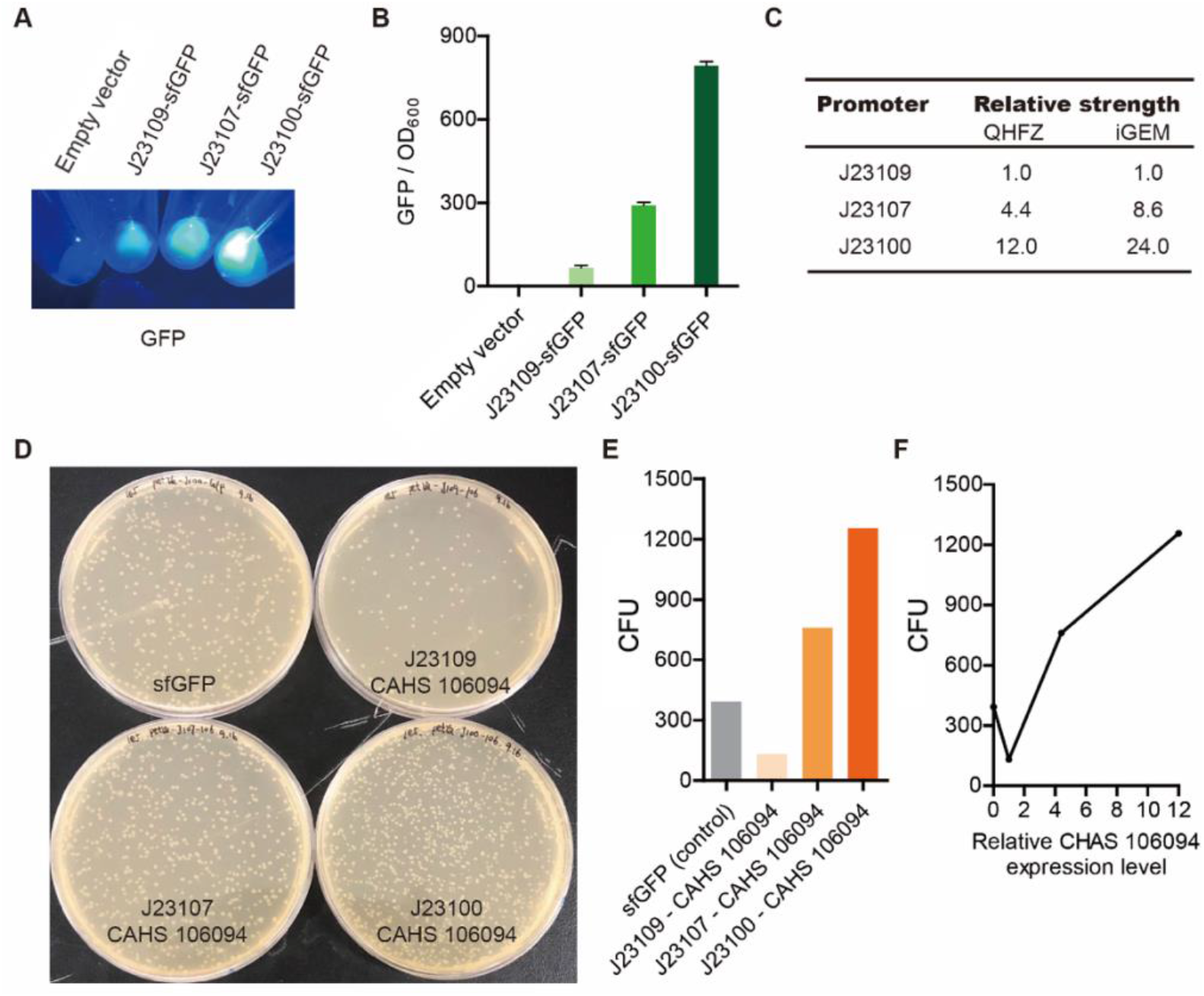
The protective effect of CAHS 106094 was enhanced after increasing its expression. (A)Visual inspection of GFP expression illustrating the relative strength of the J23109, J23107 and J23100 promoters. (B) Quantitative analysis of the assay shown in (A). (C) The relative strengths of the promoters. The strength of J23109 was defined as 1 (D) The lyophilization-recovery test of *E. coli* BL21 (DE3) strains expressing CAHS 106094 with different promoters. (E) Statistical analysis of the assay shown in (D). The data represent the means ± SD from 2 technical replicates. The CFU value is calculated with a dilution factor of 10^5^. (F) Statistical analysis of the assay shown in (D). The X-axis represents the relative expression level of CAHS 106094. The data represent the means ± SD from 2 technical replicates. The CFU value is calculated with a dilution factor of 10^5^.

We used sfGFP as a reporter during promoter selection, and the results indicated that J23109 is a weak promoter, J23107 is moderate and J23100 is strong ^[43]^. However, as each of the promoters resulted in different expression rates of the same protein, we tested them on the TDPs as well to determine whether the expression level will influence the protective effect of the TDPs and the resulting survival rate. After cloning CAHS 106094 into the vectors with each type of promoter, the strain with the J23100 promoter, which expressed the most CAHS 106094, exhibited the highest survival rate after lyophilization and room-temperature storage. This means that increased expression of TDPs (CAHS 106094) leads to hither viability of engineered strains based on *E. coli* BL21 during storage at room temperature after lyophilization.

Furthermore, we also tested two additional CAHS proteins (106094, 107838), as they shown in previous studies from TU Delft as well as Boothby to be the most efficient. Since the study will not have control over the variables if it tests different strains of TDPs at the same time, the SAHS strain was eliminated. Afterwards, CAHS expression was induced with IPTG to protect BL21 during lyophilization, resulting in an increased survival rate compared to the GFP control group. This means that the expression of CAHS proteins can to protect cells of *E. coli* BL21 after lyophilization, allowing them to be preserved at room temperature. After comparing the survival rates of strains expressing the four different CAHS proteins, it was clear that CAHS 106094 allowed the best protection under lyophilization conditions as the BL21 strain with CAHS 106094 showed the highest survival rate during room-temperature storage following lyophilization.

Result 3: Co-expression of TDPs in *E. coli* BL21 can improve the survival rate of the bacteria during lyophilization.

After the expression of CAHS in BL21 showed a positive protection effect, we hypothesized that the co-expression of multiple TDPs may lead to an improved survival rate compared to the expression of a single TDP ^[40]^. Moreover, after research and interviews, we discovered that because of the diversity of TDP protein structures and their corresponding functions, the co-expression of different TDPs is possible ^[13]^. Therefore, after predicting the complementary structures of TDPs using SwissProt, we fused CAHS 106094 to the C-terminus of SAHS 33020 and expressed the resulting fusion protein using the plasmids we previously constructed.

**Figure 3.**
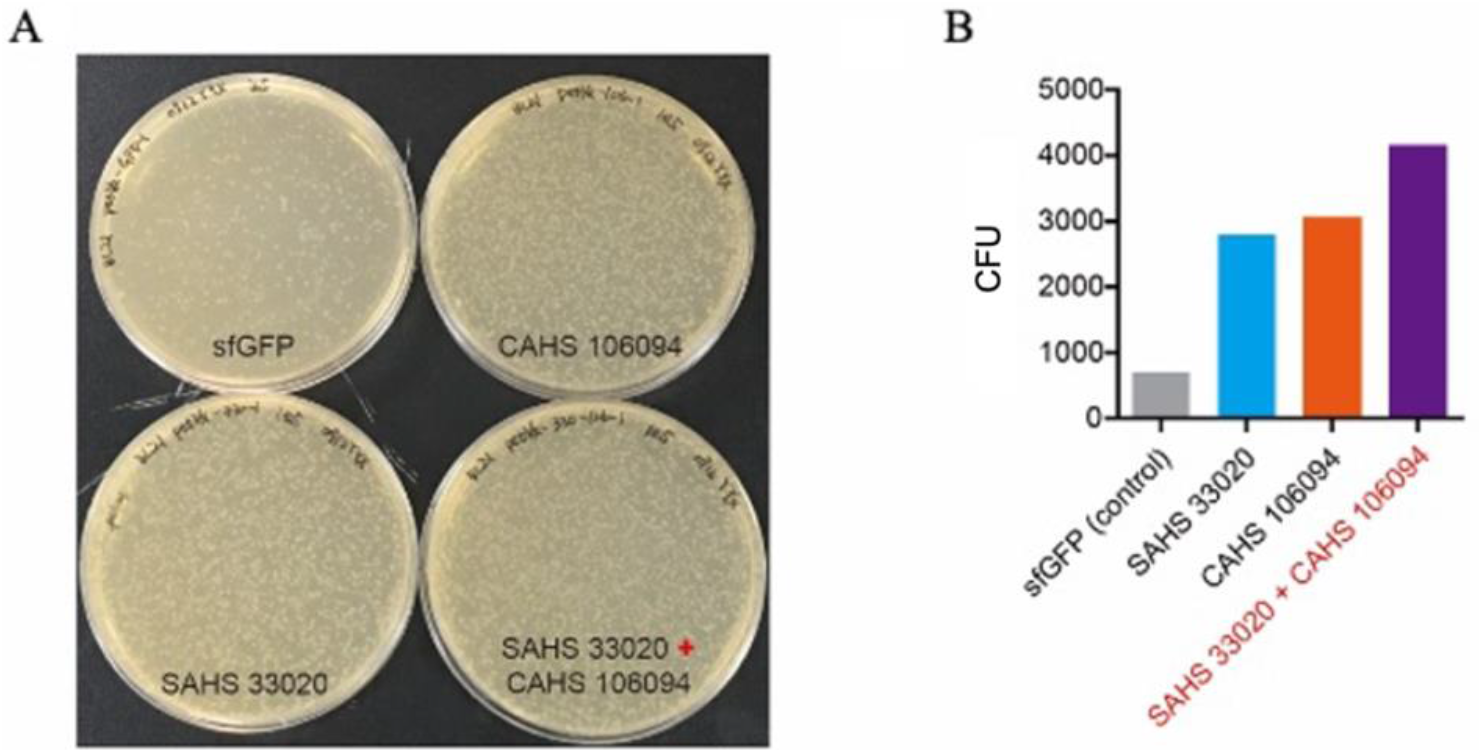
Combined expression of different TDPs (SAHS 33020 and CAHS 106094) had an improved protective effect. (A) The lyophilization-recovery test of *E. coli* BL21 (DE3) expressing a single TDP or co-expressing two different TDPs. (B) Statistical analysis of the assay shown in (A). The data represent the means ± SD from 2 technical replicates. The CFU value is calculated with a dilution factor of 10^5^.

The *E. coli* BL21 co-expressing two different TDPs had a higher survival rate after lyophilization than any of the strains expressing a single TDP. This means that the co-expression of multiple TDPs in *E. coli* BL21 can improve its survival rate after lyophilization and allow its storage at room temperature ^[3]^.

Result 4: Co-expression of different TDPs in *E. coli* BL21 allowed more bacterial cells to be preserved at room temperature for a longer period of time.

Moreover, as we extended the room temperature storage time from 2 to 12 days, the survival rate of the BL21 strain co-expressing two TDPs was higher than that of any tested strain expressing a single protein.

**Figure 4.**
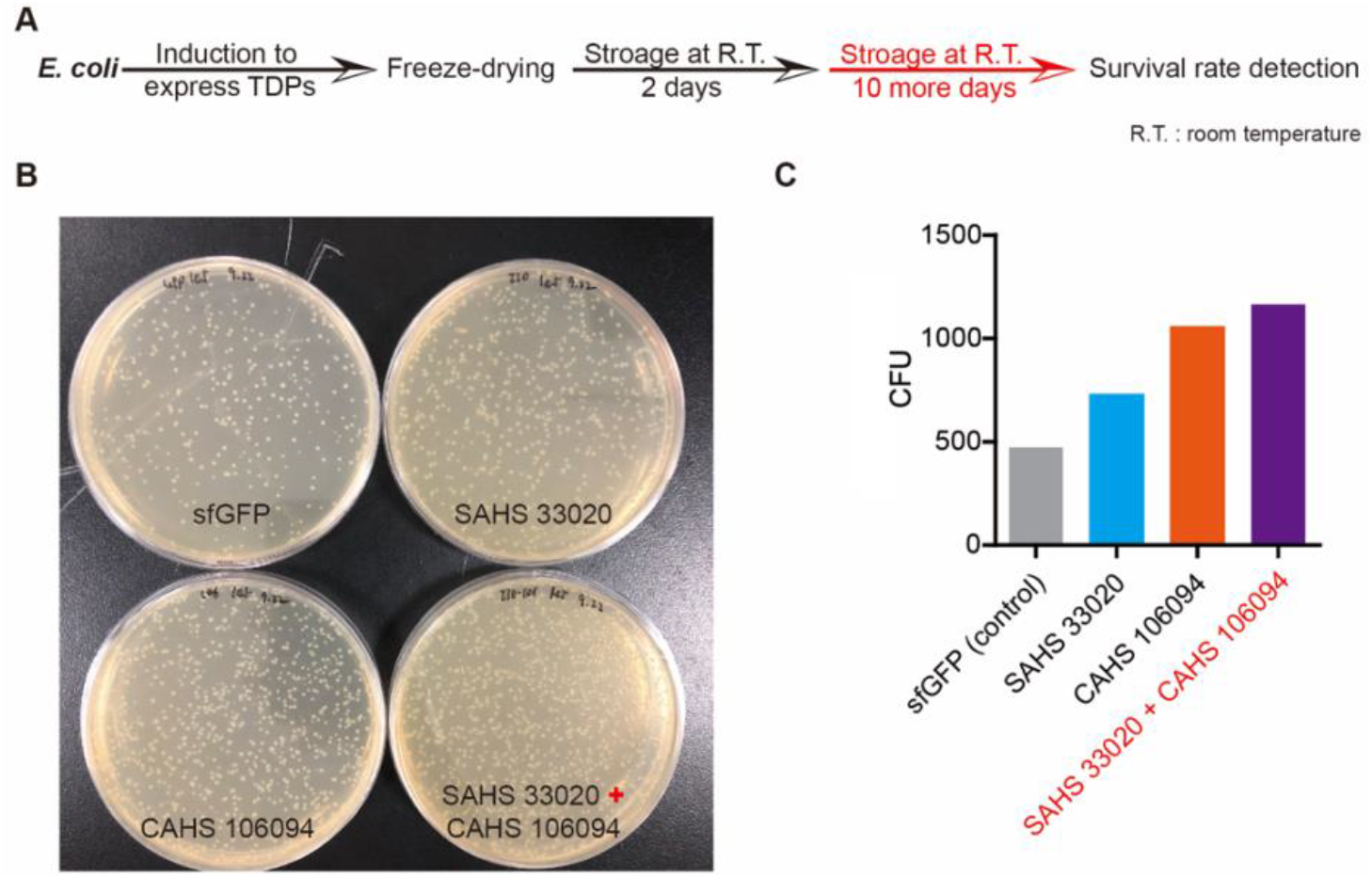
TDPs showed protective effects during long-term room temperature storage. (A) Flow chart of the experimental protocol. (B) The lyophilization-recovery test of *E. coli* BL21 (DE3) strains expressing different TDPs. (C) Statistical analysis of the assay shown in (B). The data represent the means ± SD from 2 technical replicates. The CFU value is calculated with a dilution factor of 10^5^.

Result 5: TDPs can protect different bacterial strains from loss of viability during lyophilization, allowing different bacteria to be preserved at room temperature.

After the positive results exhibited by CAHS and the co-expression of CAHS 106094 and SAHS 33020, we hypothesized that the TDPs would display the same protective effect ^[25]^ during the lyophilization of a different *E. coli* strain, as the bacteria have similar structures. Therefore, although another team failed to obtain a positive result with the DH5α strain, we utilized the pYB1a vector we received from NEFU to conduct a new set of experiments. We cloned the coding sequence of CAHS 106094 into the pYB1a vector and expressed the protein in *E. coli* DH5α. After lyophilizing the bacteria containing the modified vector and preserving them at room temperature, the results showed that CAHS 106094 was able to protect the DH5α strain against loss of viability during lyophilization and resulted in a higher survival rate at room temperature than the GFP control group.

**Figure 5.**
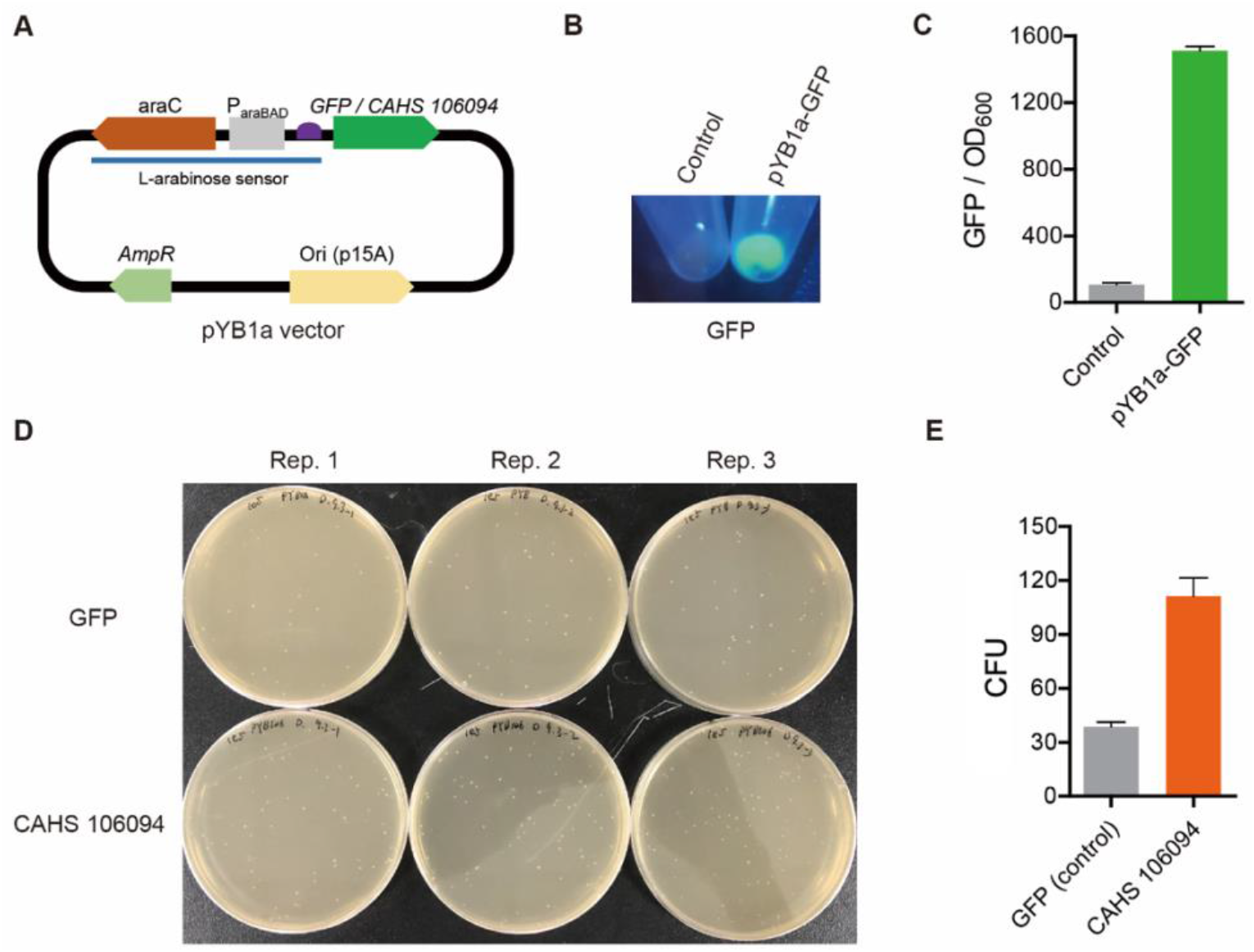
TDP (CAHS 106094) showed satisfactory modularity. (A) Schematic of the vector used to express CAHS 106094 in *E. coli* DH5α. (B) GFP as used as a reporter to confirm that the target gene in the vector can be induced with 0.2% L-arabinose. (C) A quantitative data of Fig. 5B. (D) The lyophilization-recovery test of *E. coli* DH5α strain that expressed CAHS 106094. CAHS 106094 showed protective effect against lyophilization conditions. (E) The statistical graph of Fig. 5D. Data are shown via mean ± SD from 3. The CFU value is calculated with a dilution factor of 10^5^.

After the expression of CAHS 106094 in the DH5α strain, we realized by observing the survival rate of the DH5α after lyophilization and room temperature preservation that the TDPs have a protective effect on DH5α. After editing the TNT detection vector^[36]^ (in DH5α) given to us by NEFU-China, we added the coding sequence of CAHS 106094 into the vector. After the expression of CAHS 106094, we lyophilized the DH5α strain with the TNT detecting vector and sent the sample back to NEFU-China to determine whether the original function of the TNT detector was maintained after room-temperature storage. The results showed that the TNT detection ability was still present in the bacteria and the survival rate after lyophilization as well as room temperature storage was improved.

## 4. Discussion

The main result of the study (result 1) shows that TDPs can protect *E. coli* BL21 under conditions of lyophilization ^[19, 35]^, allowing the storage of engineered strains at room temperature. A possible explanation for this effect was postulated in the water-replacement hypothesis proposed by Thomas C. Boothby et al., according to which the TDPs are able to replace the hydrogen bonds formed between water and the cellular macromolecules. Therefore, once the water molecules are removed during the lyophilization process, the hydrogen bonds between them and the cellular contents are replaced by TDPs, thereby preventing protein denaturation and membrane fusion. This result is significant because of its innovative implications. As one of the main issues in synthetic biology, the preservation of biological materials, including proteins, DNA and whole cells is extremely important. Notably, this method not only allows bacteria to be stored at room temperature, but also extends the storage time compared to previous storage methods. Moreover, as the bacteria are stored in the form of a dry powder, they can also be utilized in biomedical applications such as the shipment of pharmaceuticals very easily. Additionally, the results shown in Figure 1C were obtained using a specific lyophilization protocol, in which the bacteria were frozen for at least 2 hours at −80°C and the vacuum pump was used to dry the bacteria for at least 20h at 1 Pa. As mentioned above, the conditions of lyophilization are extreme. Due to time limitation, we only conducted the lyophilization for all TDPs a single time, and CAHS 106094 showed the best effect. Therefore, we chose CAHS 106094 as the experimental group when we prolonged the experiment, and we using pET28a and GFP as the negative control group. Moreover, the data was calculated based on the survival of colonies per 1000 cells. Hence, Figure 1C proved that the difference of survival rate between the two negative control groups and the experimental group was not significant. Thus, CAHS 106094 again showed an improvement of the bacterial survival rate ^[11]^. In conclusion, we were able to reject the possibility that GFP may have a weak protective ability, therefore confirming the validity of the control.

Result 2 shows the positive relation between TDP expression levels and the final preservation effect of the bacteria. This result is based on the idea that increased expression of protective proteins will increase their effect as long as the expression level does not exceed the equilibrium point that the cell can endure. It is significant because this means that the usage of TDPs does not only produce positive results but can also be controlled, since different expression levels enable different preservation effects. If certain applications require a specific protection level, it can be achieved by regulating the TDP expression levels using different promoters or inducers. This confirms the possibility to regulate bacterial protection by controlling TDP levels.

Result 3 and result 4 showed that the co-expression of TDPs can improve the final protection effects compared to the original individual expression of single TDPs. We hypothesized that the reason why tardigrades show high resistance to extreme environments such as extreme temperatures and vacuum in space is that there is more than one protein that protects the cells of tardigrades ^[2]^. By analyzing the protein structures, we found that SAHS 33020 and CAHS 106094 showed the most obvious difference ^[41, 44]^. The complementarity of SAHS and CAHS is due to their respective structures when desiccated. While SAHS commonly folds into a beta sheet structure, most CAHS are alpha helical proteins ^[23]^. This indicates that their combination will increase preservation levels during lyophilization as the difference in protein structure will cause the proteins to function in different preservation stages ^[4]^. The significance of this result is that the co-expression will improve the preservation effect of TDPs^[29, 44]^. Based on this, researchers can infer that combining more proteins will lead to higher survival rates of the bacteria. The individual expression of single TDPs may only protect the bacteria during one aspect of lyophilization, while using more proteins can cover more aspects of preservation. This provides a new possibility for other preservation methods that utilize resistance proteins similar to TDPs.

After testing the BL21 strain of the *E. coli*, we examined the abilities of TDPs to protect other strains of *E. coli* under lyophilization. This hypothesis was based on the understanding that different bacterial strains of the same species have similar structures and therefore may express similar proteins. The result shows that the ability of TDPs to preserve bacteria at room temperature under lyophilization is not only present in one specific type of bacteria, but is also applicable in many different strains. Moreover, as the main goal of this study was to protect engineered bacteria, this expanded usage range will allow the TDPs to protect a wide selection of engineered bacteria, increasing its usefulness. This result showed that TDPs can preserve the bacteria at room temperature while allowing them to conduct their intended functions. The applications of engineered bacteria increase in range every day, but the preservation methods limit their daily usage. We added the CAHS 106094 into an engineered bacterium that can detect TNT. After lyophilization, the TNT detection circuit was still active, confirming that engineered bacteria can be preserved a room temperature by TDPs while maintaining their original function. If TDPs are added to an engineered bacterium, it can not only achieve room temperature storage but also preserve the engineered bacteria’s original function. Therefore, the practical value of different engineered bacteria can be preserved using TDPs, allowing them to be further utilized to benefit society and the environment.

## 5. Conclusions

As the study of synthetic biology grows rapidly, the issue of storage and transportation of biological materials required for these studies has continued to grow into one of the biggest issues of the field. Therefore, the main goal of this study was to find a solution for this issue which would allow the expansion of the whole field of synthetic biology. If methods for room temperature preservation of engineered bacteria as well as other biological materials that experience storage difficulties are further explored, experiments and implementations of biological concepts would be much easier to conduct.

The main results of this study indicate that tardigrade-specific intrinsically disordered proteins (TDPs) have the ability to protect different *E. coli* strains under conditions of lyophilization and allow the bacteria to subsequently be preserved at room temperature. This conclusion is in agreement with with the hypothesis that TDPs can protect engineered bacteria under lyophilization as it shows the TDPs’ ability to protect normal bacteria ^[5]^. Moreover, we also found that CAHS 106094 displayed the most significant protective effect against loss of viability during lyophilization in *E. coli* BL21, and it can be co-expressed with SAHS 33020 to further improve the survival rate of the bacteria. Moreover, the results indicate that TDPs can protect engineered bacteria without disturbing their intended functions in some cases but can still influence the function of engineered bacteria if they disturb the original expression of proteins in the engineered bacteria. Therefore, when adapted to the specific engineering procedures of each strain, TDPs will be able to protect engineered bacteria under conditions of lyophilization, allowing room temperature preservation.

Because the use of TDPs to improve survival rates of cells under lyophilization is a very innovative concept ^[39]^, there are still many areas left to be explored. First, future researchers should combine more TDPs in the pET28a vector used in this study or other vectors to verify whether the survival rates of bacteria under lyophilization can be further improved with different combinations of co-expressed TDPs. Additionally, researchers can attempt to change the binding site of CAHS 106094 in diverse circuits like the sHYa gene circuit, so that the original function of engineered bacteria will be minimally influenced by the TDPs.

In addition to conducting lyophilization experiments, we also tested the function of tardigrade-unique DNA-associating proteins in Human HeLa cells to preserve them under UV radiation and obtained considerable positive results ^[21, 38]^. Therefore, future studies should also focus on examining the effects of TDPs on the ability of bacteria, eukaryotic cells and even isolated proteins to withstand extreme influences such as UV of high/low temperatures ^[24, 33, 36]^. These future study areas would significantly increase the application range of engineered bacteria allowing them to be utilized in increasingly extreme environments to facilitate diverse activities.

## 6. Acknowledgements

The project of this paper is based on the research created and conducted by the iGEM team QHFZ-China, led by the two authors of the paper, Yixian Yang and Zhandong Jiao. This article was advised by Xing Zhang who led the iGEM team during 2020 season and made a major contribution to the experiments, Xuan Wang who guided the experiment and proofread the article after iGEM season, and Shiqi Wang who continuously supported us in this enterprise.

This article received support from the G19 Biology Department in which the teachers of Tsinghua University High School provided a generous amount of education and gave us well-founded suggestions and guidance.

We thank Tsinghua University High School for providing us with experimental sites and funds, which has provided us with many conveniences in the process of experimentation and teaching.

It is our pleasure to be a cooperating partner of Beijing ZENO Technology Development Company. During the whole iGEM season, ZENO company gave us great support and guidance, including techniques and funds. ZENO company assisted us in conducting online meetings and offline activities. We would like to thank iGEM, we have gained a lot in this process. We are grateful to all the teams and experts that communicated and collaborate with us.

We thank Haidian District Vocational School and Tsinghua University for providing laboratory space and experimental funds throughout the experimental process.

The Beijing Taide Medicine Production Company and the YouCare Medicine Corporation also provided professional guidance and assistance for the development of our project, which we are greatly grateful for. We also wish to thank Tian Cheng Yuan Tong, BioDee, BGI, TransGen Biotech, GENEWIZ, J&K Scientific and other biotech companies for providing us ith necessary experimental instruments and materials.

## 7. Supplementary Materials

**Table S1:** Gene Sequences Used in this Project

| Name | Function | Reference |
| --- | --- | --- |
| T7 promoter | Used to express GFP, TDPs, LEA and OtsBA | [45] |
| T7 terminator | Termination of the transcription of GFP, TDPs, LEA and OtsBA | [45] |
| B0015 Terminator | Termination of the transcription of GFP, TDPs, LEA and OtsBA | [45] |
| RBS B0034 | Ribosome binding site | [45] |
| Lac operator | Repressor of gene expression | [45] |
| Promoter J23100 | Expression of GFP and TDPs | [45] |
| Promoter J23107 | Expression of GFP and TDPs | [45] |
| Promoter J23109 | Expression of GFP and TDPs | [45] |
| Pc promoter | Arabinose-inducible expression | [45] |
| PBad promoter | Expression of GFP and CAHS 106094 | [45] |
| *araC* | arabinose-inducible transcriptional activator/repressor | [45] |
| sfGFP | superfolder green fluorescent protein | [45] |
| CAHS 89226 | Cytosolic-abundant heat soluble protein<br>89226 from the tardigrade *Hypsibius dujardini* | [5, 44] |
| CAHS 94205 | Cytosolic-abundant heat soluble protein<br>94205 from *H. dujardini* | [5, 44] |
| CAHS 106094 | Cytosolic-abundant heat soluble protein<br>106094 from *H. dujardini* | [5, 44] |
| CAHS 107838 | Cytosolic-abundant heat soluble protein<br>107838 from *H. dujardini* | [5, 44] |
| SAHS 33020 | Secretory-abundant heat soluble protein<br>33020 from *H. dujardini* | [5, 44] |
| LEA | Late embryogenesis abundant protein | [27, 31, 46] |
| OtsA | trehalose-6-phosphate synthase | [6, 34, 42, 47] |
| OtsB | trehalose phosphatase | [6, 34, 42, 47] |

**Supplementary Figure 1:**
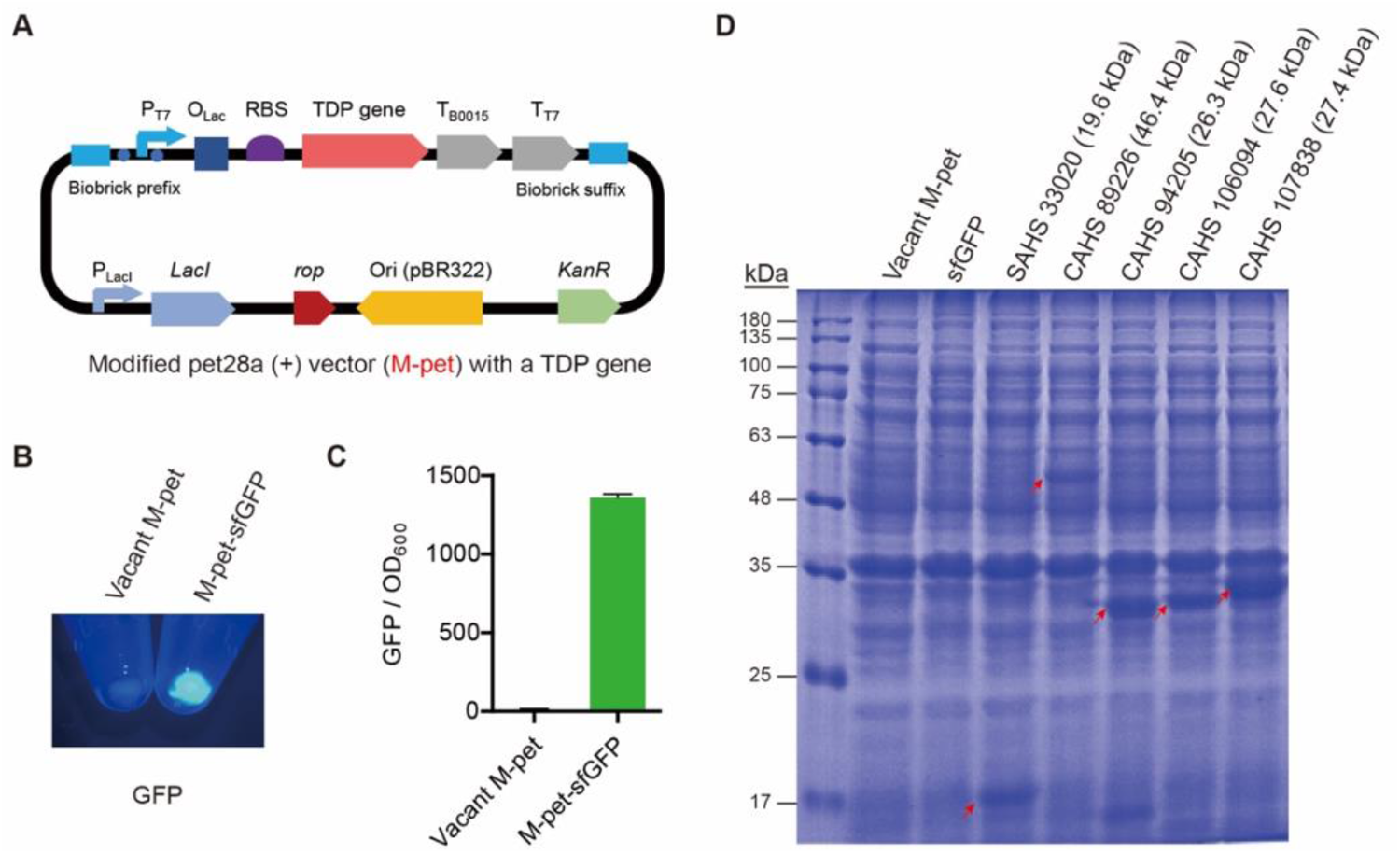
TDPs were expressed in *E. coli* BL21 (DE3) using a modified pET28a+ vector. (A) Schematic representation of the vector we used. (B) GFP was used as a reporter to confirm that the target gene in the vector can be induced with 2 mM IPTG. (C) Quantitative analysis of the data shown in (B). The data represent the means ± SD from triplicate experiments. (D) SDS-PAGE showing the expression of TDPs.

**Supplementary Figure 2:**
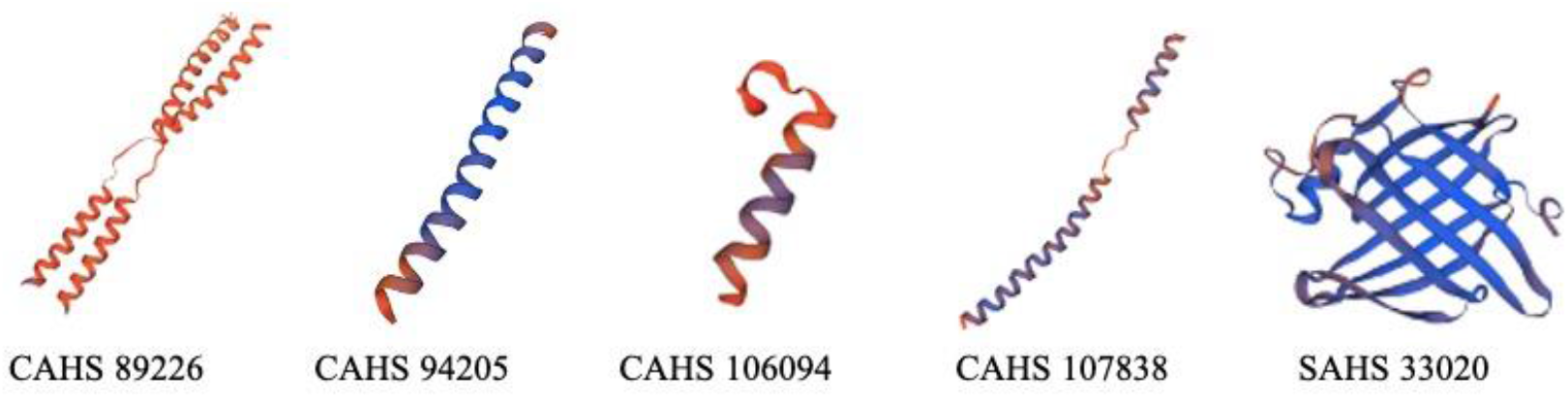
Structures of different TDPs

## 8. Materials and Methods

### 8.1 Lyophilization

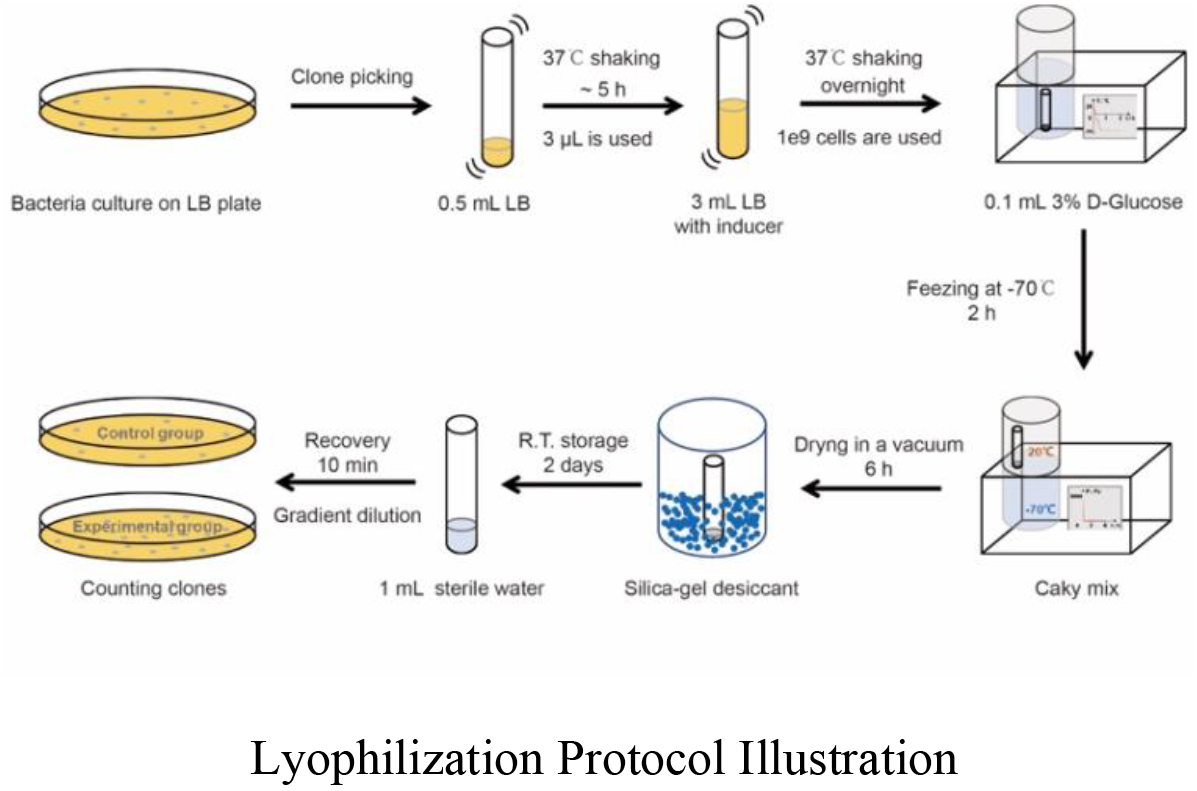

The lyophilization protocol and assessment of the results takes five days to complete. On the first day, clones that are in good condition are picked grown in 500 μL of LB medium containing appropriate antibiotics at 37°C for 5∼7 hours until the bacterial solution becomes turbid. Then, 2 mM IPTG is added into 3 mL of LB medium containing appropriate antibiotics. Afterwards, 3 μL of the bacterial seed culture is added to the IPTG-containing medium to dilute the bacteria at a ratio of 1:1000. The induced culture is then incubated in a shaker at 37 °C overnight.

On the second day, the GFP expression induced by IPTG is measured, and the experiment will be continued if the fluorescence is detectable. Then, a spectrophotometer is used to measure the OD_600_ of the bacterial solution. An OD_600_ = 1 was considered to be equivalent to 10^9^ cells. If the OD_600_ value is between 0.1 and 1, there is a linear relationship between the OD_600_ and bacterial density. The volume of the bacterial solution containing 10^9^ cells was calculated using the formula V = 100 / (OD_600_ × dilution ratio). Then, the volume containing 10^9^ cells is centrifuged at 7000× g for 3 min, the supernatant is removed, and the bacteria were resuspended in a 15 mL tube with 100 μL of a pre-refrigerated 3% glucose solution. Then, the cover of the tube is taken off and the tube is placed into the cold trap of the lyophilization machine. The compressor of the lyophilization machine is then turned on and the shake tube is allowed to freeze for 2 hours at −70°C ^[1]^. Then, the cake-like frozen solution is placed in the drying chamber of the lyophilization machine. The vacuum pump was used to dry the bacteria for 6h at 1 Pa. Lastly, the vacuum pump was turned off and the bacterial powder was placed in a sealed box filled with silica-gel, followed by room-temperature storage for 2 days.

After the room temperature storage, the survival rate of the bacteria was analyzed on the fourth day. After analysis, 1 mL of sterile water was added into the tube, vortexed for 15 seconds, and put in place at room temperature for 10 minutes. Then, the density of the bacteria was adjusted using gradient dilution and 100 μL of the bacterial solution was spread on an LB plate. The bacteria were then cultured overnight at 37°C.

On the last day, the LB plate is photographed to record the experimental results. Then, the ImageJ software is used to count the number of colonies on the LB plate.

After the iGEM season, we improved the process of lyophilization to reduce the false positive degree between two negative control groups. Accordingly, the bacteria were allowed to freeze for 2 hours at −80°C and the vacuum pump was used to dry the bacteria for 20h at 1 Pa.

### 8.2 Plasmid Construction

A total of 45 primers were designed using SnapGene software and synthesized at GENEWIZ Company. PCR was performed according to the the manuals of PrimeSTAR Max Premix (2X) (TaKaRa, R045) or 2×EasyTaq PCR SuperMix (+ dye) (Transgene, M256). The DNA fragments were separated by agarose gel electrophoresis and purified using a TIANgeo Midi Purification Kit (TIANGEN, DP209-02) or AxyPrep DNA gel Extraction Kit (AXYGEN, AP-GX-250) and stored at −20 degrees until further use.

### 8.3 Gibson Assembly

First, retrieve the previously constructed plasmids from the −20°C refrigerator. Afterwards, extract the plasmids based on their concentration and inject 5 μL of the Gibson assembly mix into the solution and add complementary volumes of sterile water to make up the total volume of the solution to 10 μL. Then, following the temperature requirements listed in the manual of the pEASY-Basic Seamless Cloning and Assembly Kit (Transgene, CU201), incubate the solution in the PCR machine to assemble the plasmids. After the plasmids are assembled, agarose gel electrophoresis is used to confirm the result, and the DNA is extracted using the TIANgeo Midi Purification Kit (TIANGEN, DP209-02) or AxyPrep DNA gel Extraction Kit (AXYGEN, AP-GX-250).

### 8.4 Endonuclease Digestion and Ligation

Retrieve plasmids that contain previously designed enzyme digestion sites (PUG, PET). The volumes of the plasmid solutions are calculate to provide 2 µg of plasmid DNA according to the concentrations of their solutions. Then, inject 5 μL of cut smart buffer, 1 μL for each of the two types of endonucleases (*Eco*RI, *Xba*I, *Spe*I, or *Pst*I), and complementary volumes of sterile water into the solution, making the total volume of the solution 50 μL. Then, incubate the solution in the PCR machine at 37°C for specific amounts of time based on the different types of plasmids. The incubated solution is then subjected to agarose gel electrophoresis and the correct band is extracted using a TIANgeo Midi Purification Kit (TIANGEN, DP209-02) or AxyPrep DNA gel Extraction Kit (AXYGEN, AP-GX-250). The extracted plasmid DNA is stored at −20°C. Then, 1 μL of T4 DNA ligase and 2 μL of T4 DNA buffer were added into the solution, and complementary amounts of digested plasmids and sterile water were added to make the final solution volume 20 μL. Then, the final solution is incubated in the PCR machine at 16°C overnight. The incubated solution is then subjected to agarose gel electrophoresis and the correct band is extracted using a TIANgeo Midi Purification Kit (TIANGEN, DP209-02) or AxyPrep DNA gel Extraction Kit (AXYGEN, AP-GX-250).

### 8.5 Plasmid extraction

After culturing the bacteria in lysogeny broth (LB) at 37°C overnight, plasmid extraction was performed following the manuals of the TIANprep Mini Plasmid Kit (TIANGEN, DP103-02) or AxyPrep Plasmid Miniprep Kit (AXYGEN, AP-MN-P-250).

### 8.6 Sequencing

Sanger sequencing was done by GENEWIZ Company (China). SnapGene software was used to proofread the sequences.

### 8.7 Transformation

Transformation was performed following the manuals of the Trans5α Chemically Competent Cells (Transgene, CD201) or Trans1-T1 Phage Resistant Chemically Competent Cells (Transgene, CD501).

### 8.8 SDS-PAGE

Bacteria are cultured overnight and their OD_600_ is measured. The 3∼5×10^9^ number of bacterial cells is lysed in 100∼200 μL 4% SDS for 5 min at room temperature, and denatured for 10 min at 95°C. Afterwards, 6× loading dye is added. The SDS-PAGE was carried out on 12% precast polyacrylamide gel. Coomassie brilliant blue stain is used to observe the bands.

